# Impacts of batch effects on the performance of machine learning classifiers across multiple studies

**DOI:** 10.64898/2026.06.24.734352

**Authors:** Preston Raab, W. Evan Johnson, Stephen R. Piccolo

## Abstract

Precision medicine relies on accurate and generalizable predictions for patients across the spectrum of human diversity. Because capturing biological heterogeneity requires large sample sizes, researchers must often aggregate data from several experimental batches or independent studies. This integration allows for greater statistical power and diversity than a single study could provide, while avoiding the costs of generating massive new -omics datasets. Predictive models trained on these aggregated data are theoretically better equipped to detect subtle patterns that generalize to new data. However, this potential is frequently undermined by "batch effects"—systematic technical artifacts that can bias model training to predict experimental batches and shadow meaningful biological conditions. Models trained on data with batch effects can exhibit substantially degraded performance when applied to data from new batches. Statistical adjustment methods can mitigate these artifacts while preserving biological signals. To ensure these adjustments actually facilitate generalization, we emphasize the use of external, independent cohorts for rigorous validation. This chapter examines how batch effects impact predictions and compares various adjustment methods.

## Classification in Precision Medicine

Classification—the task of assigning observations, or patients, to predefined categories based on their characteristics—is vital to biomedical research and the implementation of precision medicine. Indeed, precision medicine has been defined as “the tailoring of medical treatment to the individual characteristics of each patient…to classify individuals into subpopulations that differ in their susceptibility to a particular disease or their response to a specific treatment” [1, 2]. For instance, treatment for cancer patients can be made more precise by classifying tumors into molecular subtype groups. In fact, several classifiers have been approved and implemented clinically to predict personalized recurrence risk for breast cancer and guide chemotherapy decisions [3, 4].

Before a classifier can be used for prediction, it must be trained on labeled data: a collection of samples where the correct class, or label, is known. A classifier cannot be tested using the same data it was trained on; classifiers can easily achieve perfect performance on seen data. Instead, evaluation requires a held out set of labeled data that is not used for training. A well-trained classifier will generalize to new, unseen data, which could be new patients in need of prediction. To train a classifier that generalizes, the training set must contain a sufficient number of samples to represent each class well, with good coverage over the many possible individuals that could belong to each class.

Biomedical classification tasks can utilize diverse data types, including protein abundance measurements, genomic sequences, clinical variables, and imaging data. Each of these data modalities provides complementary information about biological systems and disease mechanisms. No modality is immune to batch effects, though routine clinical measurements, such as weight, are resilient [5]. Batch effects arise from differences in experimental protocols, equipment type and condition, reagent lots, environmental conditions, human error, and other technical factors that vary between studies or even within studies over time. Generally, greater complexity and subjectivity in measurement procedures imply greater susceptibility to batch effects. For our purposes we will focus on gene expression. Gene expression data provide an excellent context for understanding how batch effects can be modeled and removed, as the technical variation introduced by measurement technologies is well-characterized and substantial. Insights from gene expression are extensible to several other biological modalities and likely apply to high-dimensional data more broadly.

### Gene Expression Data

RNA measurement technologies quantify the abundance of RNA transcripts in biological samples. These measurements provide a snapshot of cellular activity, reflecting which genes are actively expressed under specific conditions, disease states, or in response to treatments [6]. Gene expression data enable the identification of disease-associated pathways, the development of diagnostic and prognostic biomarkers, and the discovery of therapeutic targets [3, 7]. Public repositories, like the Gene Expression Omnibus (GEO), offer hundreds of thousands of studies and millions of samples. For each molecular sample, data submitters frequently provide relevant clinical variables, enabling associations between molecular signatures and clinical phenotypes [8]. These repositories provide researchers free access to integrate labeled gene expression data across multiple studies. This integration offers substantial benefits over single-study analysis: meta-analyses can reveal consistent patterns across diverse populations, increased sample sizes improve robustness to noisy data, and independent validation cohorts provide evidence of generalizability to new populations.

Batch effects pose challenges for multi-study integration due to the sensitivity of transcriptomic technologies to technical variation. These technical artifacts are often comparable to or larger than the biological signals of interest, making them a primary concern when combining datasets. They appear in gene expression data as systematic shifts in expression levels, differences in variance structure, and even alterations in the relationships among genes [9]. Batch effects can also manifest as changes in distributional shape for features shared between datasets. For gene expression data, this difficulty arises when combining legacy microarray values with modern RNA sequencing (RNA-seq) values. Across samples, log-transformed microarray measurements of expression intensity tend to be more normally distributed (bell-shaped), with continuous values. In contrast, RNA-seq technologies measure discrete transcript counts, resulting in heavily right-skewed distributions. Direct comparisons between the two are impossible without intervention. Fortunately, the right-skewed RNA-seq distributions for genes with large count numbers can be modified using log-based transformations to yield distributional shapes comparable with microarray data. Shifts in expression levels and differences in variance structure remain to be addressed.

### Batch Correction for Gene Expression

Batch effect correction algorithms, statistical adjustment methods, or *adjusters* for brevity, aim to remove technical artifacts while preserving biological signal. Adjusters typically employ one of two strategies: modeling and subtracting the technical batch effects, or modeling the biological subspace and removing extraneous variation through projection. This choice has implications for signal preservation. If a biology-focused model fails to capture some component of biological variation, then that signal will be discarded. Similarly, if an adjuster that models batch effects attributes too much variation to the batch, that variation will be removed. The effectiveness of adjusters depends on the validity of the assumptions made. These modeling assumptions vary by data modality, as demonstrated in the following comparison.

Among the subtypes of gene expression measurements, two are dominant: bulk, which measures total expression in a large sample of cells, and single-cell, which measures the expression of individual cells. For bulk data, a common adjustment approach is ComBat. ComBat addresses batch effects by modeling batch-specific shifts and scaling factors for each gene [10]. Once modeled, these effects are subtracted or divided out. The transformation that ComBat applies is a robust, regularized (defined later) linear model: each expression value for a particular gene is shifted and scaled uniformly. This approach works well for bulk gene expression data, where values vary continuously due to the random proportions of cell types represented in the sample.

Single-cell data requires, and allows for, a more specialized adjustment. Single-cell gene expression is characterized by distinct expression signatures for each cell type, with internal variation caused by various cell states at measurement time. If bulk data spans a continent of continuous variation, single-cell data occupies islands of specialized differentiation. Single-cell adjusters may assume that cells of the same type will exhibit similar expression patterns even across batches. By finding "nearest neighbors"—samples with the most similar expression profiles between batches—adjusters such as Harmony or Seurat v3 can identify how to move cross-batch neighbors closer together to bring the batches into alignment [11–13]. Because these methods find neighbors using many genes and apply sample-specific transformations, the adjustment of single-cell data is typically nonlinear. These batch correction methods model the biological space, then eliminate the batch differences.

ComBat and the single-cell methods mentioned also differ by mathematical “space”, or set of variables they use. ComBat is a *feature-space* correction method: it attempts to fix the original data and return new values for each sample and gene. Harmony and Seurat are *latent-space* or *embedding-based* alignment methods: they represent each sample using a smaller set of derived variables which represent simple ways the data can vary. These new variables are called latent, hidden, or embedded, because they were not among the original genes yet hold most of the information. By performing adjustments in this lower-dimensional space, these methods reduce computational load but are unable to offer corrected values for the original genes.

Latent space adjustments share many similarities with models developed for domain adaptation. Both attempt to remove information about the batch or domain of the data in a latent space. Domain adaptation focuses on finding domain-invariant representations of the data in layers of deep neural networks through training [14]. This requires large amounts of data and is not the focus of this chapter, but the idea of a domain-invariant representation is a useful framework. It is not generally necessary to preserve the original data distribution for classification. If a transformation of the data preserves the essential information while removing potential batch effects, the transformation would act similarly to batch-aware correction techniques for the purposes of classifying the data.

### Confounding

Batch effects can mimic or mask real differences in expression levels and covariance structures among batches [15]. Because not all differences among datasets are technical artifacts, not all differences should be removed. For example, consider two studies having the same balance of positive and negative cases but having opposite proportions of female and male subjects. Since models trained on imbalanced data generalize poorly for underrepresented classes, it would be ideal to combine these datasets with reciprocal imbalances [16]. However, if the studies are suspected to vary technically, it will be difficult to determine whether observed batch differences are technical artifacts that should be removed, or true differences due to the sex imbalance between datasets. We say that the batch effect is "confounded" with sex—the effect of the batch on gene expression is mathematically entangled with the effect of the population imbalance [17].

Confounding can often be mitigated if the values of the secondary variables are known and recorded. For example, ComBat, when provided with additional sample metadata, can temporarily isolate gene expression from metadata associations, remove the remaining batch effects, and then reintroduce the metadata associations. However, if these other variables are unknown—due to poor recording or unknown population differences—addressing confounding becomes substantially more difficult. In the multi-study context, these additional variables are rarely consistently recorded [18]. One variable might be deemed important to one study but omitted in another. These unshared variables typically cannot be used for merging datasets. Even when variables are shared, inconsistent naming and non-standardized values add practical hurdles to combining datasets.

In extreme cases, datasets are perfectly confounded, meaning the batch variable and the target class carry identical information. This occurs if, for instance, one study contains only "Normal" samples and another contains only "Pneumonia" samples. In a two-study merge, a classifier cannot mathematically distinguish between the biological pathology and the batch-specific signatures. In fact, whenever the target class is fully confounded with another variable—such as all pneumonia samples coming from children and all normal samples coming from adults—models can learn the wrong patterns and fail to generalize [17, 19]. While specialized methods like LIGER use matrix factorization to separate shared biological features from dataset-specific ones, perfect confounding often represents a fundamental limit to what can be safely integrated [20].

A separate but related challenge arises when the batches—the distinct non-biological sources of variation within a dataset—are latent or are unknown. While this is less of a concern when combining separate datasets, it is a frequent issue within single studies. Methods such as surrogate variable analysis (SVA) identify and adjust for these hidden batches by extracting latent variables that capture unwanted variation without prior knowledge of the source [21, 22].

### Single-Patient Data

In precision medicine, single-patient data also poses a major obstacle to batch correction. Single-patient data processing is vital for translating molecular assays into clinical settings, as patient samples in clinical settings are typically collected in small numbers, often one at a time. However, many correction techniques rely on several samples to characterize the distribution of the new batch. Without this reference distribution, it is statistically difficult to know if a single sample is an outlier or merely dominated by technical artifacts.

If possible, it is best to process several samples at the same time or use recent samples collected using identical protocols to define the distribution of the new data. If this is not feasible, transformations may be applied in an attempt to shift individual samples into a batch-independent space, including within-sample rank-based methods and deep autoencoders for latent variable extraction [23, 24]. Unfortunately, the performance of within-sample transformations are often inferior to methods that utilize information from multiple samples. A simple example of this effect is seen in Figure 2, where adding an additional ranking over samples (Rank Twice, described in Table 4) improves the performance of feature rankings (Rank Features). To understand the impact of these and other adjustments on classification, we will first describe the modern classification landscape.

### Machine Learning Classifiers for Gene Expression Data

Classification has evolved from traditional statistical methods, such as logistic regression and linear discriminant analysis, toward modern machine learning approaches, such as random forests [25], neural networks [26], and gradient boosted trees [27]. This direction increasingly favors flexible methods that can capture complex, non-linear patterns in high-dimensional data. Modern techniques often use iterative and stochastic (random) training, improving the model in small steps to minimize classification error. Simple, rigid models possess a high inductive bias, meaning they rely on strong assumptions about how inputs relate to labels. This bias provides protection against overfitting to idiosyncrasies of the training data that do not reflect the distribution of the classes [28]. More complex machine learning models are less constrained and can fit more complicated relationships. This flexibility can lead to highly variable model parameters in the face of random sampling.

For bulk gene expression classification, comprehensive reviews and benchmark studies consistently highlight several highly effective algorithm families [3, 29]. Support vector machines (SVMs) are particularly designed for the high-dimensional, few-sample challenges inherent to gene-expression data [30]. SVMs identify decision boundaries by maximizing the margin between classes, relying strictly on the most informative samples, or support vectors. Ensemble tree-based methods, including random forests and XGBoost, also deliver strong performance [6, 31]. By aggregating predictions from many simpler decision trees, these algorithms manage complex feature interactions and limit overfitting [25, 27]. For scenarios where ample sample sizes are available, neural networks—ranging from multi-layer perceptrons to advanced architectures—provide excellent classification capabilities [32, 33]. Regularized linear models, despite their simplicity, also demonstrate good performance on binary classification tasks [6].

### Regularization

Many machine learning algorithms use some form of regularization, or assumptions of simple relationships, which constrain the models. This can improve generalization, or the ability to classify unseen data. Complex models rely on regularization to avoid overfitting. Even for simple models that fit one or two parameters per feature, high-dimensional gene expression data cause models to fit thousands of parameters. Logistic regressors succumb to the noise of tens of thousands of genes when training, using small contributions from many genes to fit the training data exactly. By penalizing the model for each gene with a non-zero weight, the number of genes used by the model can be limited [34]. Limiting the number of features is called feature selection. Feature selection, done manually or incorporated into models, is important whenever the number of features far exceeds the number of samples used for training. It encodes the idea that only a few features are important for predicting the label.

Automatic feature selection in weighted models can be performed using L1 regularization [35]. The L1 penalty drives coefficients of irrelevant features to exactly zero, creating models that use few features. This results in interpretable and computationally efficient models. Feature selection can also be done post-classification by examining which features were important for classifying the training or validation data. Strategies such as permutation importance can perform this calculation for any classifier type [36]. In the context of precision medicine, feature selection finds minimal diagnostic signatures. This can reduce thousands of genes to a small panel suitable for cost-effective clinical assays. However, this utility depends entirely on selecting genes that represent genuine disease biology rather than technical artifacts. A minimal signature derived from batch-confounded data will fail catastrophically when deployed in a clinical setting with different technical conditions.

Generalization to new settings also requires resilience to noisy data. Fitting training data exactly is equivalent to fitting noise that will not be found in other settings. Overcoming noise requires regularization, such as the L2 penalty. The L2 penalty penalizes large coefficients, which encodes the idea that genes or features with low variation should not have a large effect on classification. The idea that features should not be weighted too strongly is a general idea that is widely applicable to all data types. Neural networks, and other models that utilize weights can, and often do, use L1 and L2 penalties. Linear regression with the L1 penalty is known as LASSO [37]. Regression with the L2 penalty is known as ridge regression [38]. The term ElasticNet is used when both penalties are incorporated into the model [34].

### Ensemble Classifiers

Regularization can also be incorporated by training many simple learners into a more complex ensemble model. Since each simple learner is constrained in its ability to fit the data, this imposes a bias that adds regularity to the larger model. These ensemble methods are robust to the high-dimensional, noisy nature of gene expression data because they aggregate predictions across small models trained on random subsets of samples and features. This averaging effect reduces sensitivity to outliers. One popular ensemble model is the random forest [31]. Random forests construct ensembles of decision trees, where each tree is trained on a bootstrap sample of the data and a random subset of features [25]. For gene expression data, this means each tree sees a different combination of genes and samples, preventing small pattern deviations from dominating the model. The final prediction aggregates votes across all trees, providing robustness to noise and the ability to capture complex interactions between genes. Random forests also provide measures of feature importance, which can aid in biological interpretation by identifying which genes contribute most to classification decisions. A similar model, XGBoost, builds trees sequentially to correct errors from previous iterations, often achieving excellent performance on structured data [27, 39]. It benefits from the same regularization methods as random forests.

### Evaluating Batch Adjustment

Batch effects can introduce systematic biases that classifiers learn to exploit, leading to inflated performance on merged training data but poor generalization. Effective batch correction removes these technical artifacts, allowing classifiers to focus on biological signals and improving external-study performance. External-study performance serves as a good indicator of both biological preservation and batch reduction because it directly tests whether a classifier can generalize to independent datasets with potentially different technical characteristics [40]. High performance on unseen external data suggests that batch effects have been successfully removed while biological signal has been preserved.

Testing against unseen batches, without adjustment, usually results in failure. Exceptions occur if the unseen batch is remarkably similar to a batch used in training. In theory, if datasets representing the full joint spectrum of batch effects and biological variability were assembled, a classifier could then generalize easily to any new data. In most cases, the data to be predicted on must be adjusted to match the training set, without modification of the data used in training. This homogenizes the data used as input to the classifier. Any datasets merged to form the training sets should also be adjusted, to prevent "shortcut learning" on technical differences unrelated to biology [41].

Principal component analysis (PCA) and other visualization methods can help identify batch effects but do not directly measure their impact on classifier performance [42]. If points are colored or shaped by label, separation between batches indicates that biological signal needed for classification is present in the most variable genes. After adjustment, PCA often shows visual overlap among batches. Though PCA might show overlap in the two most variable directions, it cannot confirm the absence of separation in other directions. Neighbor mixing metrics identify the proportions of nearest neighbors that share a batch or a metadata label. Ideally, samples are well mixed by batch and poorly mixed by label. This gives some insight into whether a classifier may perform well across datasets, but does not guarantee superior classification outside of nearest-neighbor based classifiers. Additional tools to visualize batch effects can be useful. BatchQC provides interactive software for evaluating sample and batch effects with multiple diagnostic approaches including PCA, heatmaps, dendrograms, and statistical metrics [43]. These visualizations help researchers assess whether batch effects have been successfully removed while preserving biological structure, though these methods, especially those relying on visual separation, are inconclusive.

### Interaction Effects Between Adjusters and Classifiers

Different batch correction methods preserve or remove different aspects of the data structure, which can interact with how different classifiers learn decision boundaries. Some adjusters use the same modeling assumptions as particular classifiers, which can lead to improved relative performance [44]. Studies which evaluated many transformations and classifiers have consistently found that adjuster performance depends on the classifier model, with some top-performing adjusters showing minimal interactions (Table 1). Conversely, studies which evaluated fewer combinations often did not find these effects. This suggests that interactions are real but rare, requiring a high number of classifier-adjuster pairings to detect. We therefore conducted a reanalysis of TB transcriptomic data to determine if these are stochastic coincidences or technically grounded synergies. We identify several distinct pairings where performance deviates significantly from the mean. Meaningful interactions are technically feasible and should be considered during pipeline selection.

**Table 1:**
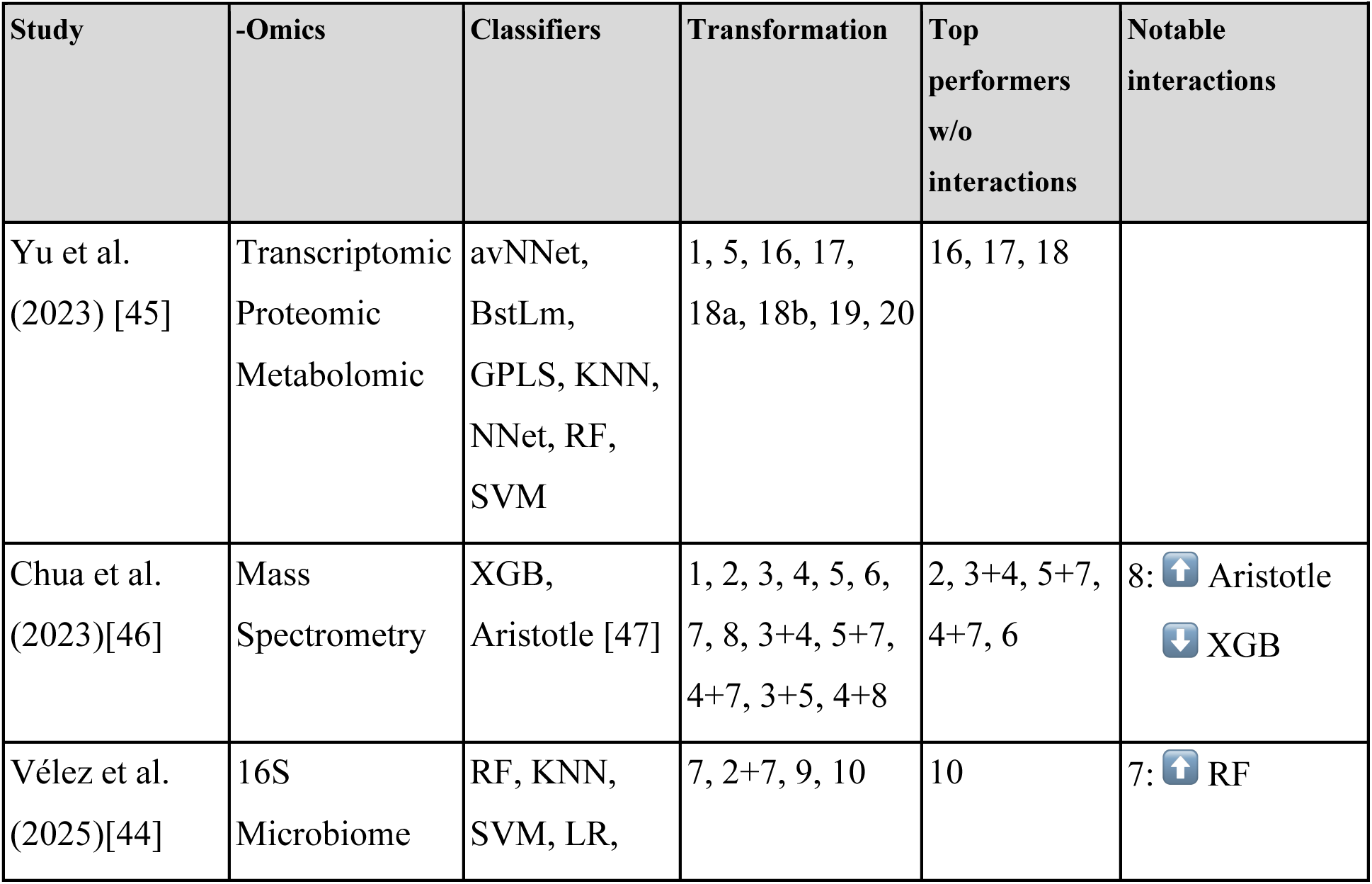
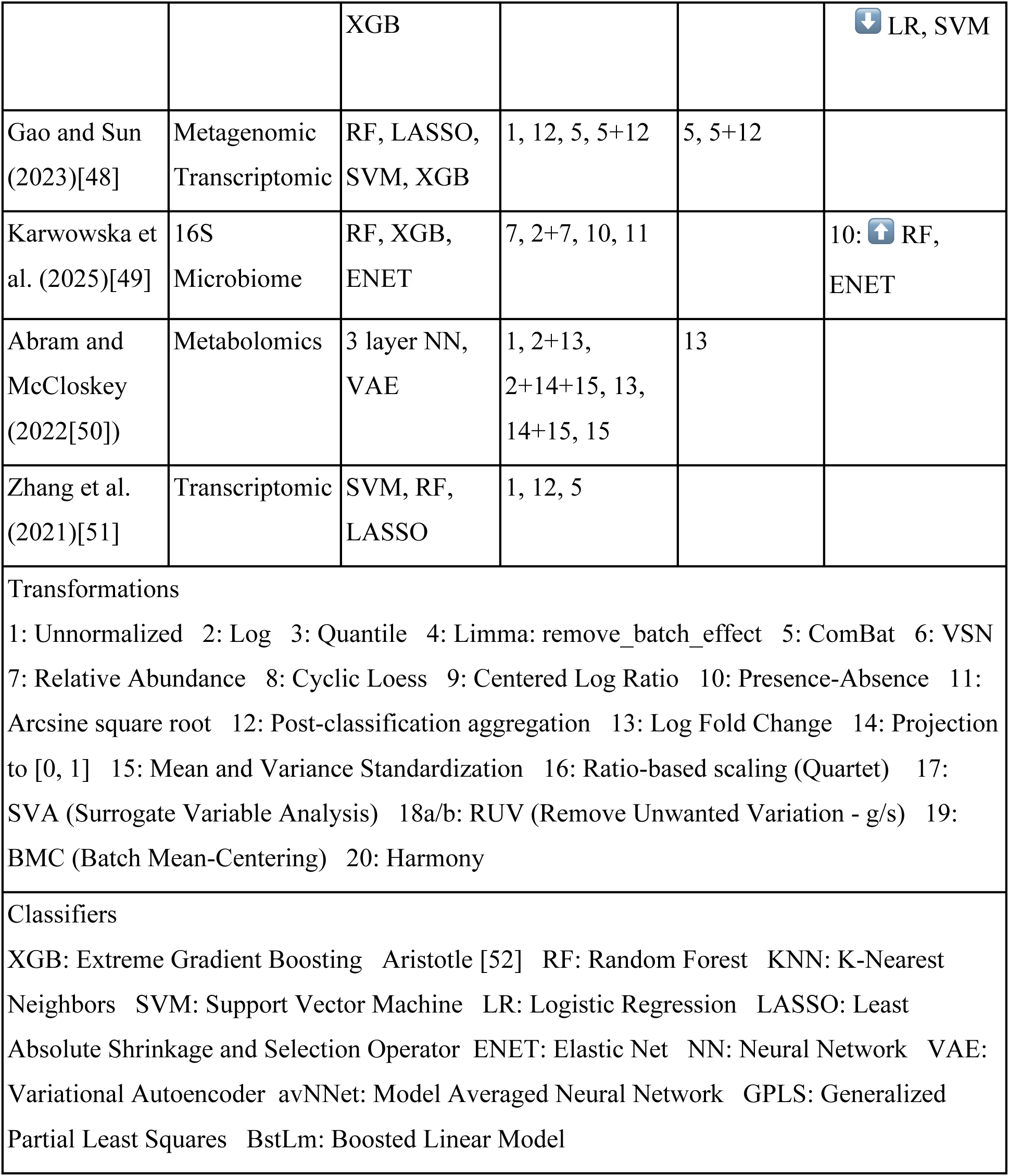
Studies Analyzing the Performance of Multiple Transformations and Classifiers in Omics Research. Summary of published studies evaluating how data transformations interact with machine learning classifiers across diverse omics modalities. Columns indicate the data type, classifiers tested, transformations evaluated (numbered below), and which methods performed best overall versus those showing classifier-specific performance shifts (interactions). Arrows denote relative changes in performance for specific classifier–transformation pairings. Across studies, interactions are most frequently observed when a large number of transformation–classifier combinations are evaluated, suggesting that such effects are real but may be overlooked in smaller comparisons.

### The Link Between Adjustment and Classifier Performance

As indicated in Table 1, Vélez et al. found that when classifying microbiome data, centered log-ratio (CLR), log-transformed relative abundance, and presence–absence representations consistently outperform raw relative abundance (RA) for logistic regression and SVMs with radial-basis (Gaussian) kernels, but not for Random Forest or XGBoost classifiers [44]. This pattern can be explained by how these transformations alter the geometry of compositional data. CLR and log-based transformations map data from the simplex (a space where features are relative proportions constrained to sum to 1, forcing artificial correlations) to an approximately Euclidean space in which log-ratio relationships become linear and distances are more meaningful, aligning with the inductive biases of logistic regression and distance-based kernels. In contrast, tree-based methods such as random forests and gradient boosting rely on threshold-based feature splits and are largely insensitive to global geometric structure, resulting in weaker dependence on the transformation. Presence–absence transformations, while discarding abundance information, similarly reduce dominance effects from highly abundant taxa, which benefit the models sensitive to feature scaling. The KNN classifier generally underperformed without strong interaction effects, which the authors attribute to sparsity in the data. The results of Vélez et al. illustrate a broader principle: transformations that reshape the geometry of the data can either align with or violate the assumptions of a classifier, producing systematic differences in performance.

To show that similar interaction effects arise in transcriptomic data, and to visualize the impact of batch effects on classifier performance, we reanalyze tuberculosis data assembled by Zhang et al., also analyzed by Gao and Sun [48, 51]. This collection of datasets deliberately included studies spanning pediatric, adolescent, and adult populations, collected across multiple continents using both microarray and RNA-seq platforms. Although the original studies encompassed diverse research objectives—including longitudinal progression risk and clinical diagnostics—all datasets in this analysis were harmonized to a standardized binary classification task: distinguishing active tuberculosis from latent infection. These diverse cohorts, with substantial technical and biological variation, test the usefulness of batch correction methods for broad precision medicine applications. Study details are summarized in Table 2.

**Table 2:**
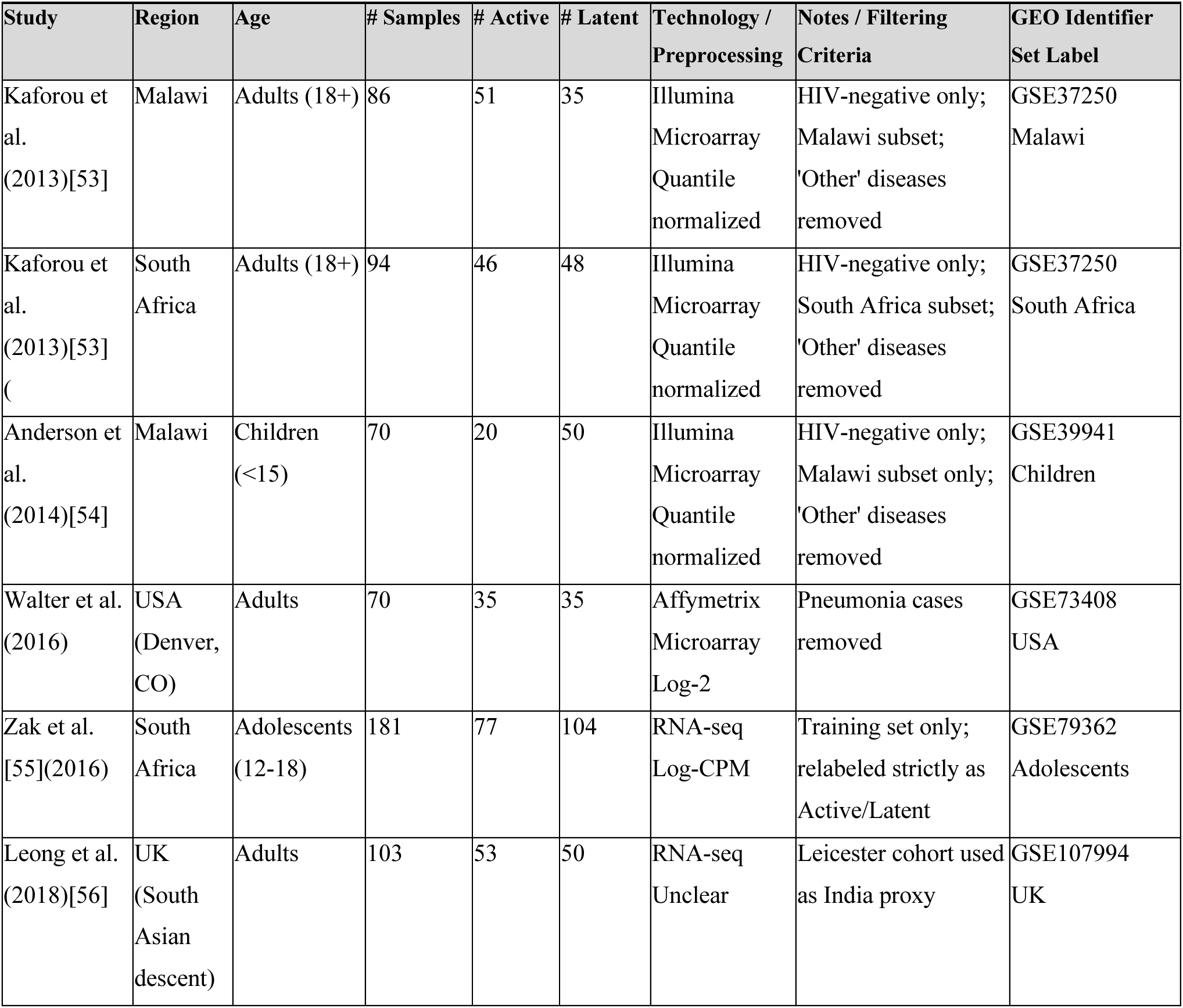
Tuberculosis Datasets Reanalyzed. This table outlines the geographical origin, demographic diversity (age), sample sizes, and sequencing technologies for the six cohorts assembled by Zhang et al. to test batch effect adjustment methods.

| Study | Region | Age | # Samples | # Active | # Latent | Technology / Preprocessing | Notes / Filtering Criteria | GEO Identifier Set Label |
| --- | --- | --- | --- | --- | --- | --- | --- | --- |
| Kaforou et al. (2013)[53] | Malawi | Adults (18+) | 86 | 51 | 35 | Illumina Microarray Quantile normalized | HIV-negative only; Malawi subset; 'Other' diseases removed | GSE37250 Malawi |
| Kaforou et al. (2013)[53] | South Africa | Adults (18+) | 94 | 46 | 48 | Illumina Microarray Quantile normalized | HIV-negative only; South Africa subset; 'Other' diseases removed | GSE37250 South Africa |
| Anderson et al. (2014)[54] | Malawi | Children (<15) | 70 | 20 | 50 | Illumina Microarray Quantile normalized | HIV-negative only; Malawi subset only; 'Other' diseases removed | GSE39941 Children |
| Walter et al. (2016) | USA (Denver, CO) | Adults | 70 | 35 | 35 | Affymetrix Microarray Log-2 | Pneumonia cases removed | GSE73408 USA |
| Zak et al. [55](2016) | South Africa | Adolescents (12-18) | 181 | 77 | 104 | RNA-seq Log-CPM | Training set only; relabeled strictly as Active/Latent | GSE79362 Adolescents |
| Leong et al. (2018)[56] | UK (South Asian descent) | Adults | 103 | 53 | 50 | RNA-seq Unclear | Leicester cohort used as India proxy | GSE107994 UK |

## Methods

We tested seven classifiers (described in Table 3) across the six tuberculosis gene expression datasets on a cross-study classification task. The classifiers were trained on the latent/active label on datasets combined using nineteen adjusters. The adjusters were chosen for potential to interact with the classifiers and reduce batch effects in bulk gene expression data. The classifiers were then tested on a separate dataset that was adjusted to match the training sets. Sun and Gau note that performance can be improved by adjusting the training sets to the test set before training the classifier. While they noted improved performance on test sets seen by the adjusters before training, they did not test the trained models on independent test data. Here, we perform the standard test of a classifier’s ability to generalize to unseen data. This was repeated for all combinations of training and testing datasets.

**Table 3:** Classifiers.

| Method | Libraries / Functions | Parameters |
| --- | --- | --- |
| Support Vector Machine (SVM) | e1071::svm<br>e1071::tune | cost: Tuned from a sequence of values<br>kernel="linear": Linear decision boundary |
| ElasticNet (ENET) | glmnet::cv.glmnet | family="binomial": Logistic classification<br>alpha=0.5: 50/50 blend of L1 and L2 penalties |
| Random Forest (RF) | ranger::ranger | Uses default ranger hyperparameters |
| Neural Network (NN) | nnet::nnet | One hidden layer, limited to 10 hidden nodes.<br>decay=0.01: L2 regularization |
| K-Nearest Neighbors (KNN) | class::knn | k: Tuned from {3, 5, 7, 9, 11} |
| XGBoost (XGB) | xgboost::xgb.cv<br>xgboost::xgb.train | max_depth=3: Limits tree depth<br>eta=0.1: Learning rate<br>subsample, colsample_bytree =0.8: Uses 80% of samples and features per tree |
| Regularized Discriminant Analysis (RDA) | klaR::rda | gamma=0.3: Covariance shrinkage<br>lambda=0.6: Diagonal shrinkage |

Performance was assessed using the Matthews correlation coefficient (MCC) [57]. Because MCC is a symmetric metric that incorporates all four confusion matrix categories, it is impossible to achieve a high MCC if a classifier has poor sensitivity, specificity, positive predictive value, or negative predictive value. Consequently, a high MCC guarantees strong performance across all these individual metrics, providing a reliable lower bound for overall model evaluation [58]. MCC ranges from 1 (perfect) to 0 (random) to −1 (perfectly inverted). Performance can also be measured using the area under the receiver operating curve (AUROC or AUC). This metric is useful for choosing an informative model before selecting a decision threshold, but it is not sensitive to positive or negative predictive value [59].

Ten batch adjustment methods—selected for their variety in potential interactions with classifiers, rather than their overall performance—are described in Table 4. For the datasets not already log-transformed, the formula *New Data = log2(1 + Data - Global Batch Minimum)* was applied before other adjustments. After this transformation, constant additions in expression represent multiplicative fold changes. We note that ComBat-seq is designed specifically for RNA-seq count data and uses a negative binomial model that fits the distribution well [60]. However, for compatibility with the microarray data, we use standard ComBat and a log transformation.

**Table 4:**
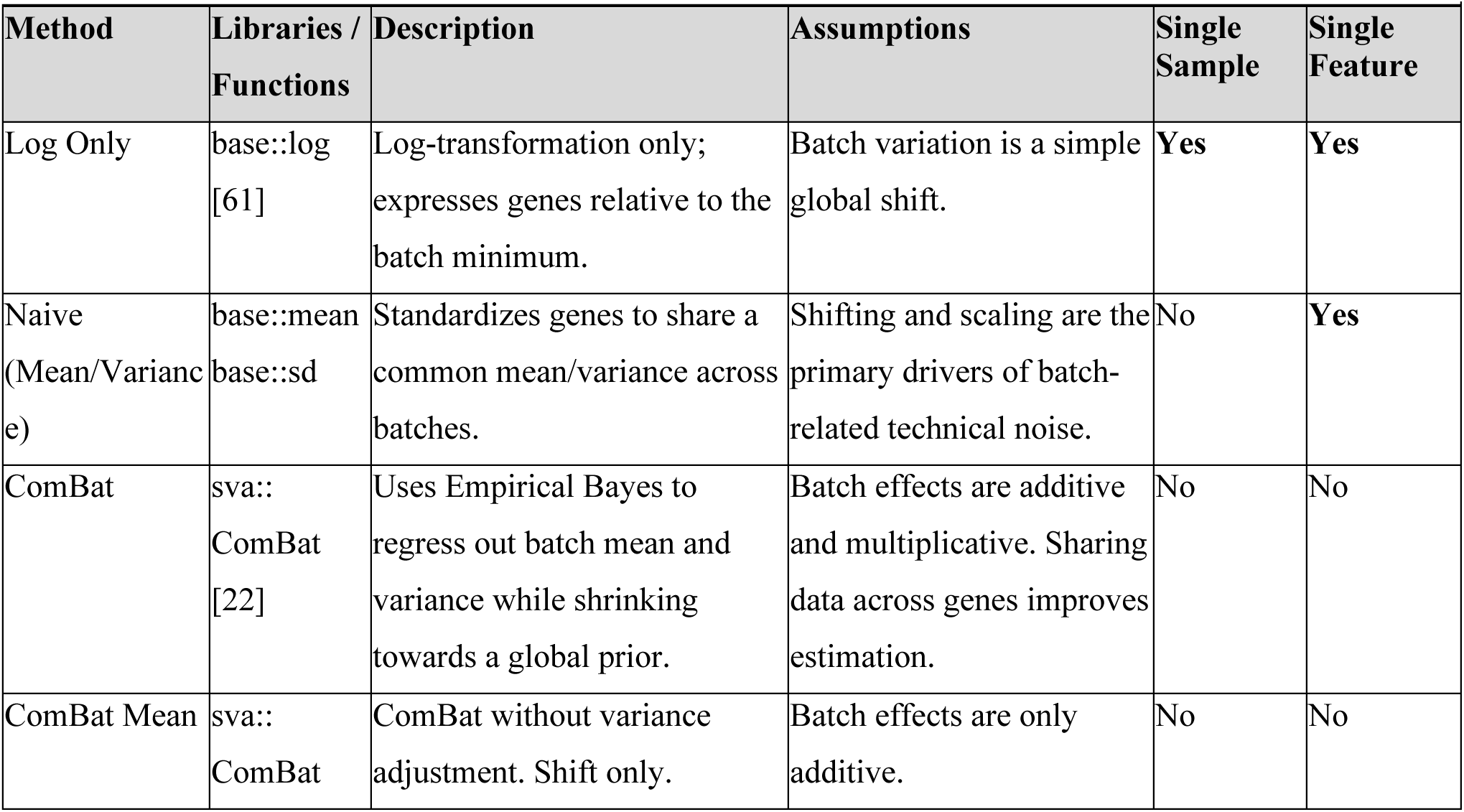
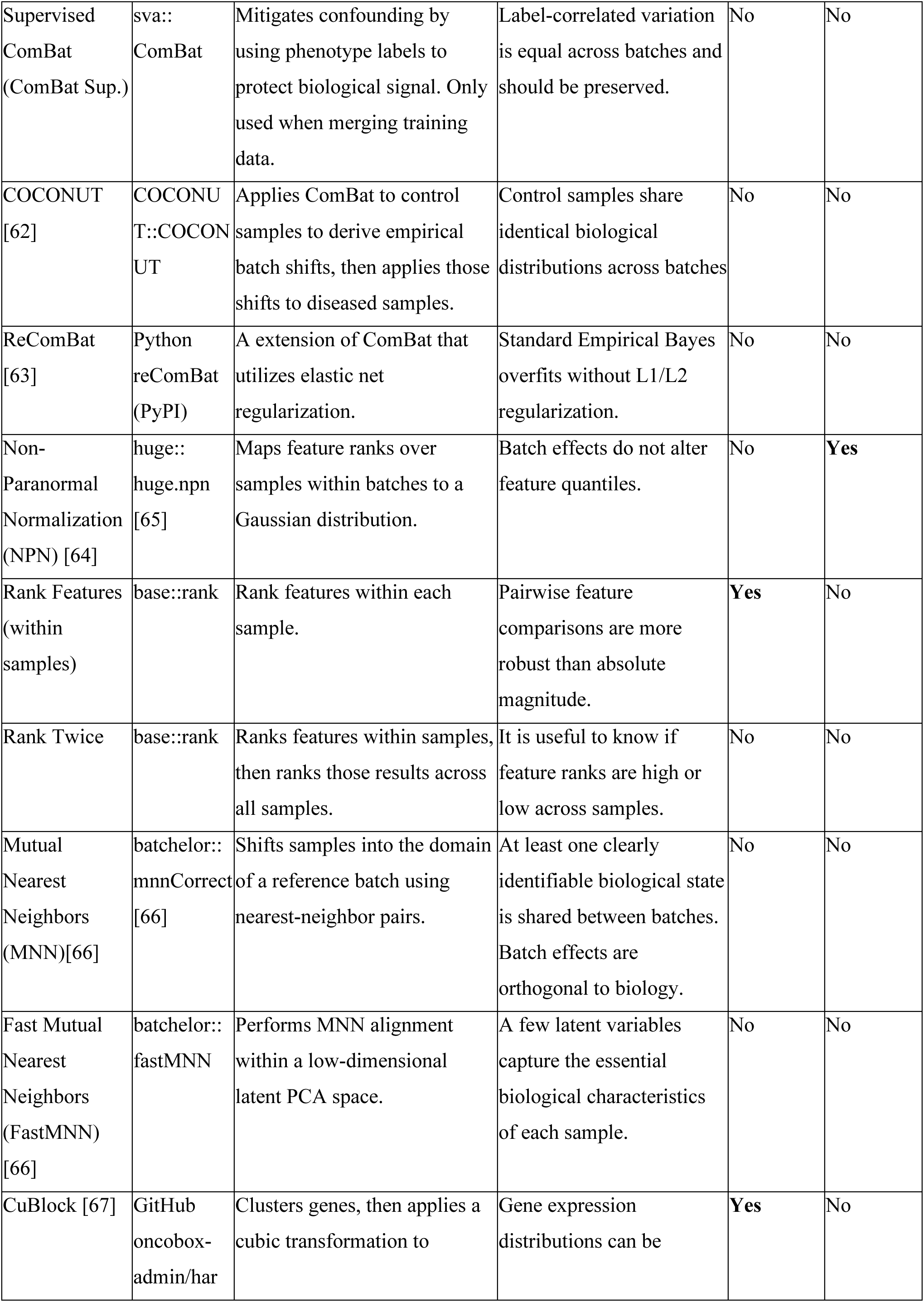
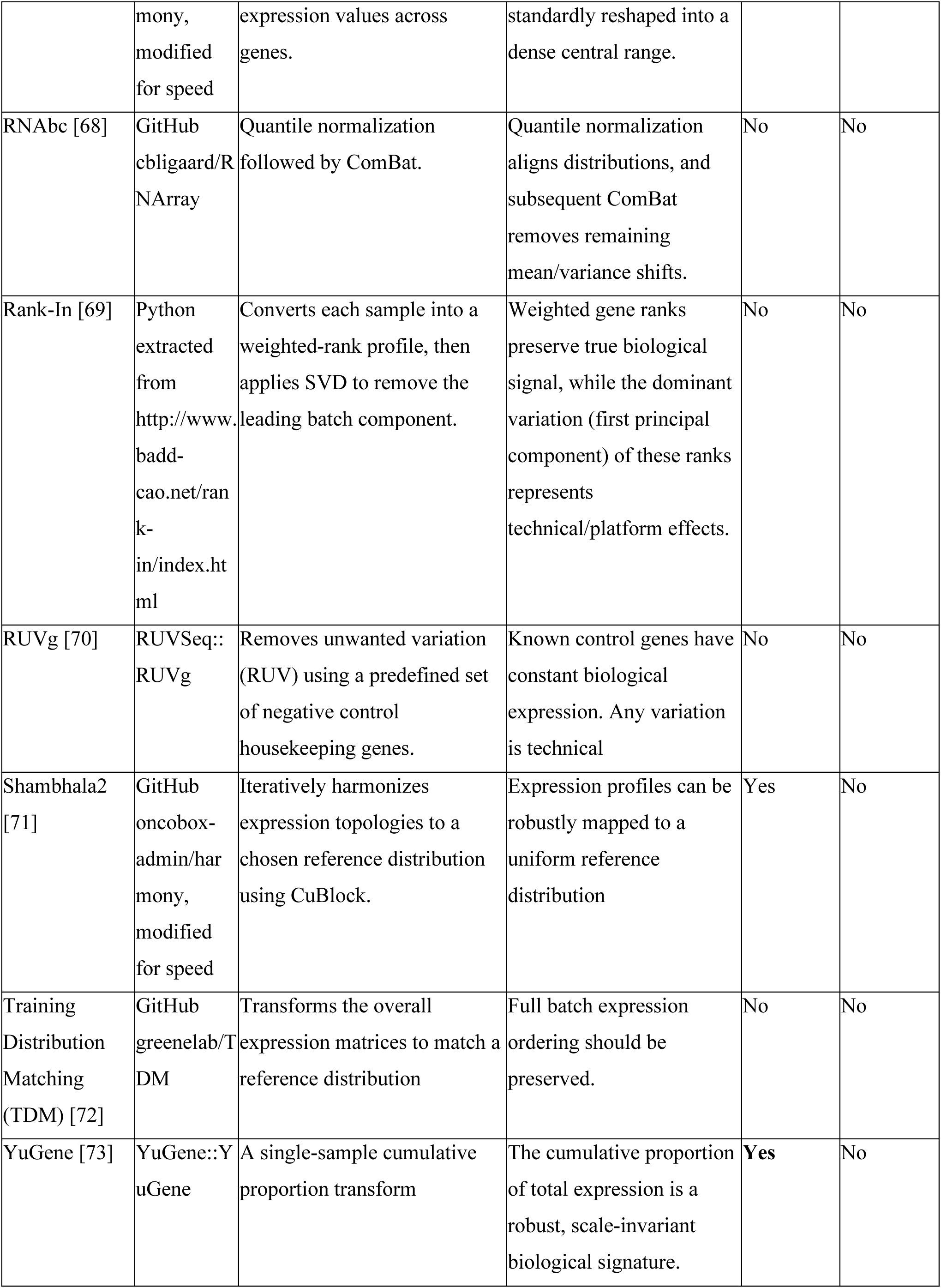
Transformations and Batch Effect Correction Methods. *Methods, Descriptions, Assumptions, and Libraries Used*

## Results

Figure 1 shows the distribution of MCC scores across training pairs and test sets, focusing primarily on classifier performance. Classifiers that can generalize from the training sets to the test sets have high (rightward) distributed MCC scores. Color indicates adjuster groups based on overall performance and interaction profiles. While interactions between classifiers and adjusters are more noticeable in Figure 2, Figure 1 shows a stark interaction between KNN and supervised ComBat, shown in light green. The L1/L2 regularization of ElasticNet enabled robust performance, with the highest overall median and fewest random results near 0 MCC. The performance of Regularized Discriminant Analysis was highly variable. Most classifier MCC distribution exhibit a multimodal structure, which results from the varying difficulties of the discrete test studies.

**Figure 1:**
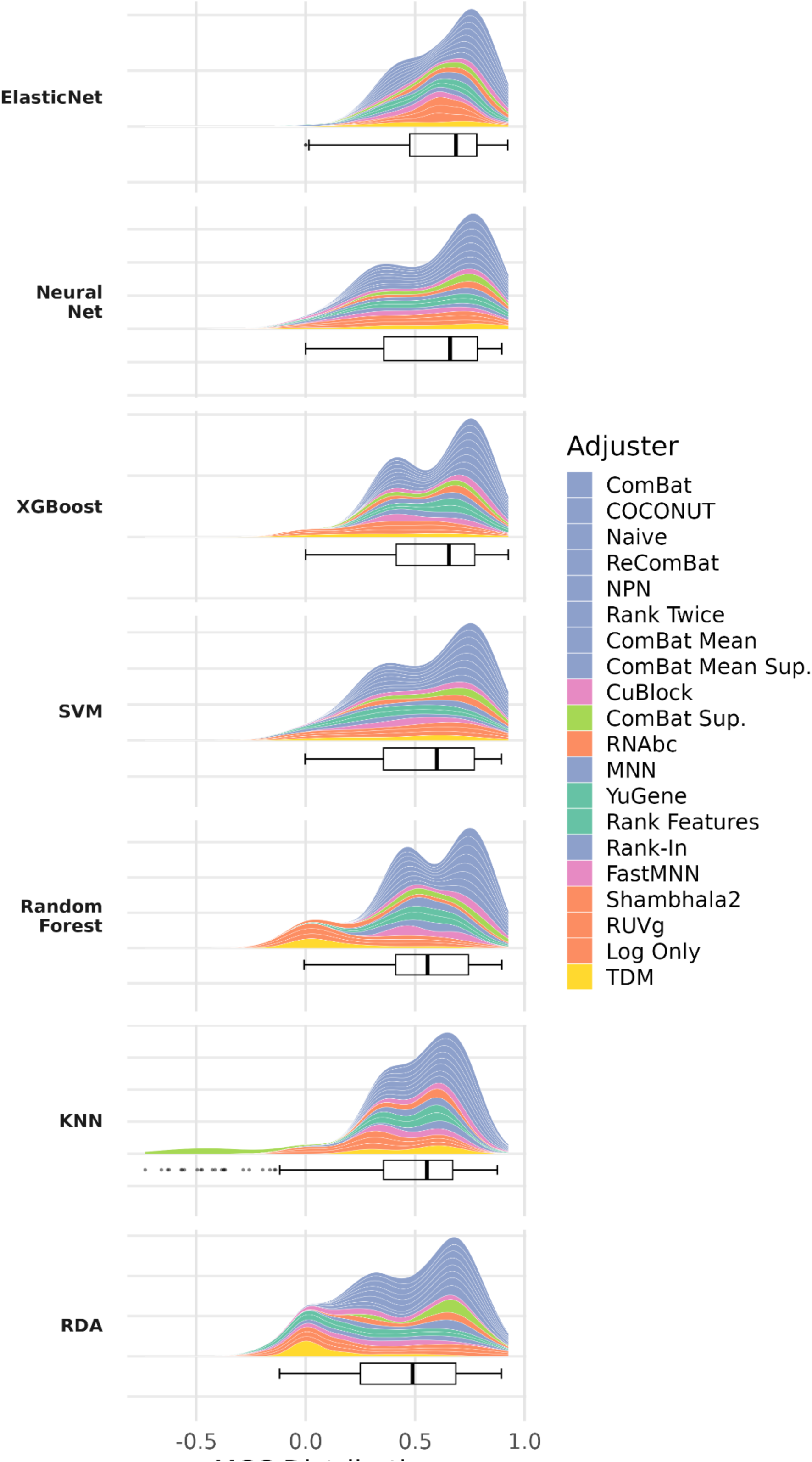
Classifier performance on cross-study tasks. Classifiers are labeled on the left. Classifiers were trained on two datasets merged using an adjuster, and tested on a third heldout dataset, adjusted to the joint training set. The full distributions of MCC performance (over adjusters, training pairs and test sets) are shown as stacked Kernel Density Estimation (KDE) plots. Adjusters are stacked vertically according to their mean performance, and colored by similarity in performance. Boxplots are shown beneath the KDE plots. Note: the maximum and minimums are clearly visible on the box plots, while the KDE plots spread density around each point.

Figure 2 shows how each adjuster ranks when paired with a classifier, highlighting interactions where an adjuster performs better or worse than typical. Most adjusters performed consistently. The top five adjusters (Combat, COCONUT, ReComBat, Naive, and Rank Twice) interacted little with any classifier.

**Figure 2:**
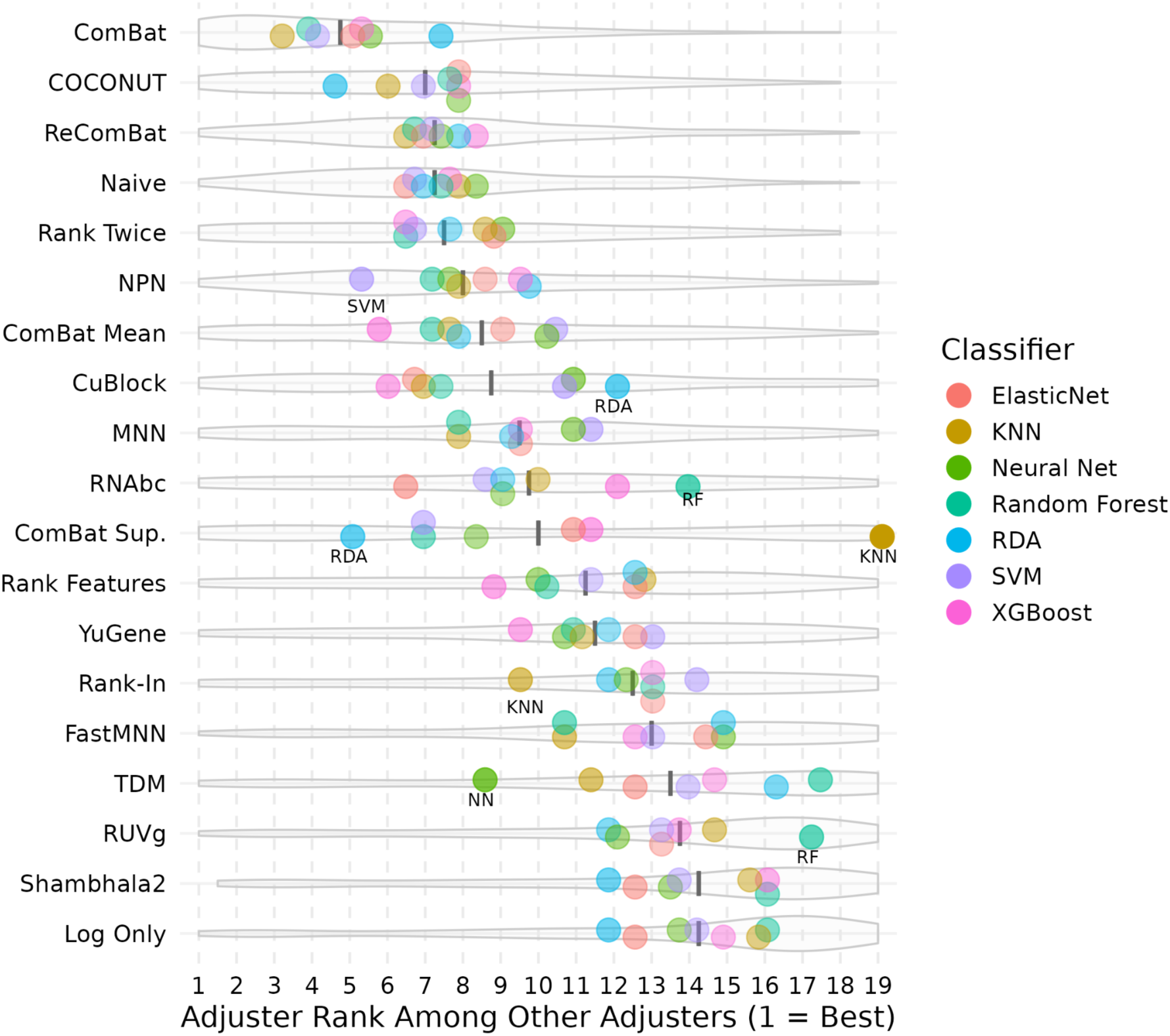
Stability of Batch Adjustment Performance Across Classifiers. Horizontal violin plots illustrate the distribution of performance rankings for various adjusters across multiple cross-study validation scenarios. Methods are ordered by their pseudomedian performance, shown as grey vertical bars, with higher-performing adjusters positioned toward the top. The x-axis represents the relative rank compared to other methods, where a rank of 1 (leftmost) indicates optimal performance. Gray violins represent the density of ranks across all scenarios. Colored points denote the pseudomedian rank achieved on specific classifiers (e.g., Random Forest, XGBoost) when using the corresponding adjustment method. Outlier classifiers that deviate significantly from the method’s central tendency are opaque and labeled for clarity. Results for Linear Regression are omitted—random performance regardless of adjuster resulted in uniform rankings that misrepresent adjuster central tendencies.

Supervised ComBat adjustment exhibits a particularly severe interaction with KNN, while matching unsupervised ComBat’s performance with RDA (see also Figure 1). These interactions arise from a mismatch between ComBat’s assumptions and the data structure, resulting in a flawed adjustment.

ComBat with covariates assumes equal effect sizes across batches: a change in class should result in the same change in gene expression for each dataset. When this assumption fails, the single pooled class effect 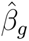 estimated across all studies is a compromise that fits no individual study exactly. Each study *i* deviates from this pooled estimate by 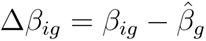, where *β_ig_* is that study’s true per-gene class effect. ComBat treats the unexplained within-batch variation—which contains Δ*β_ig_*—as technical noise. The batch scale factor decomposes as:

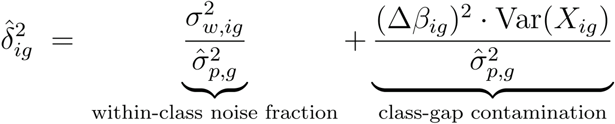

Where 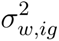 is the pooled within-class variance for study *i* and Var(*X_ig_*) = *p_ig_*(1 - *p_ig_*) is the class-proportion variance in that study. This equation holds for the raw scale estimate; empirical Bayes shrinkage contributes negligibly. The denominator 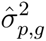 is the cross-batch weighted average of the same per-batch quantities:

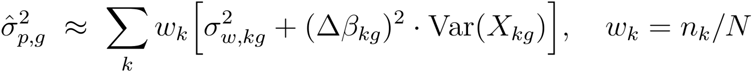

Because 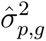 absorbs the variance from every batch, 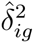 is a relative quantity—batch *i*’s variance relative to the training average—not an absolute noise measure. Batch *i* falls below the threshold 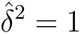 whenever its own variance is below that average, which can happen by two independent routes: if its class-effect deviation |Δ*β_ig_*| is small relative to peers, or its within-class noise 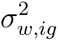 is low. Figure 3A shows the per-study distribution of 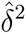, with the threshold at 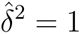 marked; Figure 3B shows the fraction of each study’s 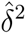 attributable to the class-gap contamination term.

**Figure 3:**
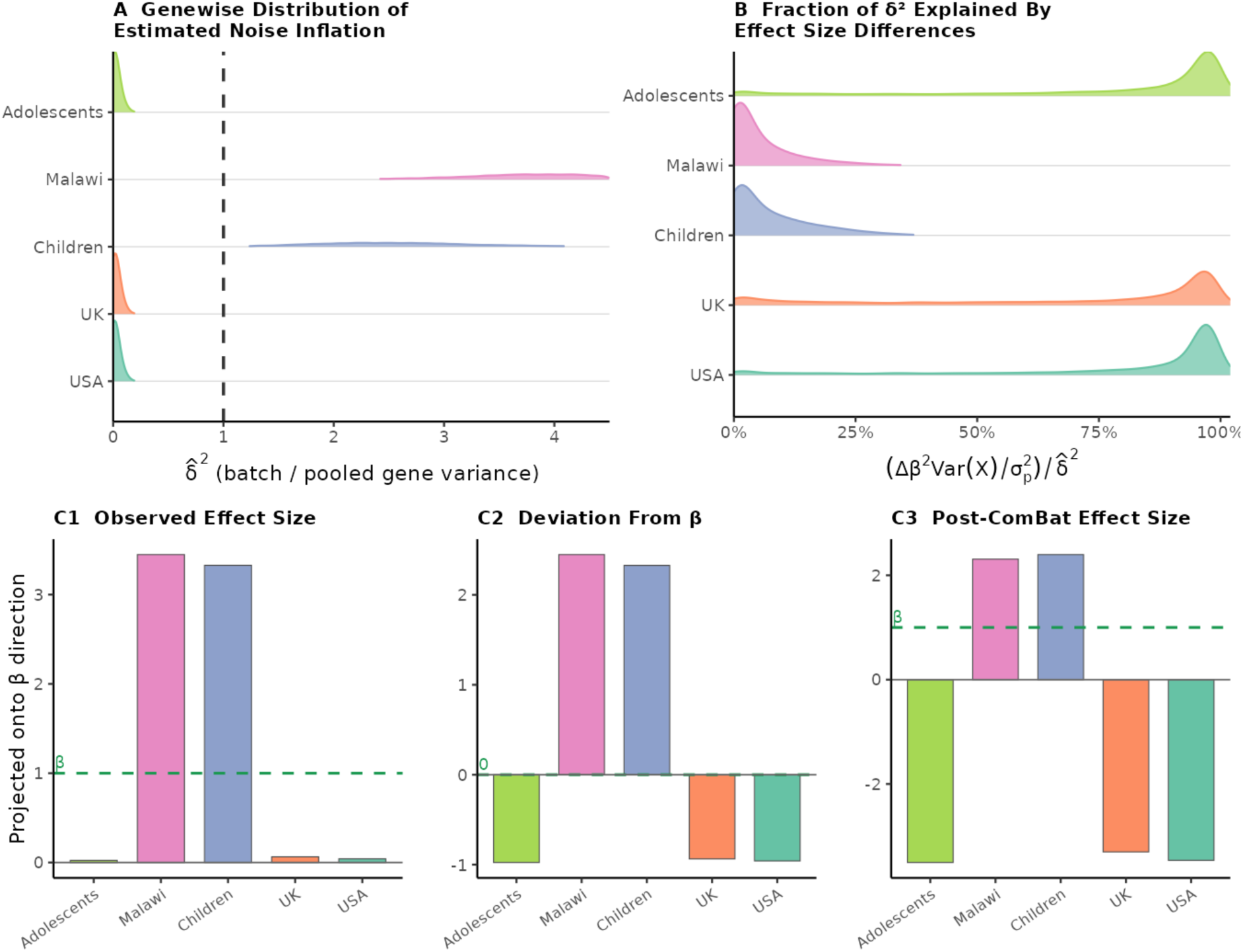
Supervised ComBat Reverses Class Effects When Effect Sizes Vary Between Batches. The behaviour of ComBat with 5 training studies is shown, similar effects are observed with fewer training sets. A: Per-study distribution of the estimated relative noise scale factor, 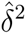, per gene, with the threshold at 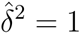 marked. B: the fraction of each study’s 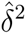 attributable to the class-gap contamination term. Effect size differences explain most of the estimated noise in low-variance cohorts, while high-variance cohorts contribute more intrinsic noise. C1: The difference between class means, projected onto the unit 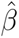 vector, for each dataset. C2: The effect sizes as deviations from 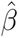, the pooled effect. C3: ComBat does not conserve class effects on a batch or pooled level when the data does not meet its assumptions.

When 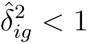, ComBat amplifies rather than shrinks within-batch residuals. Because those residuals contain Δ*β_ig_*(*x_j_* - *p_ig_*), amplification drives the corrected class axis away from the consensus 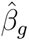; for a study whose local class effect is below the pooled estimate, the per-gene class effect can be inverted if 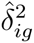 is sufficiently small. Panels C1–C3 of Figure 3 trace the overall class effect as a projection onto the 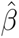 direction. The Adolescent, UK and USA cohorts all experience a strong class effect inversion. This inversion dominates the overall class effect of the adjusted data; for this example of training sets, the cosine similarity between the original and post-correction 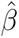 is a striking −0.75. What is surprising, then, is not that KNN fails with supervised ComBat-corrected training data, but that the other classifiers—RDA in particular—retain any ability to generalize.

MNN and FastMNN both perform well when combined with the KNN classifier. The neighborhood adjustments that the MNN methods make are easily recognized by the KNN classifier, but do not transfer as well to the other classifiers.

High-performing adjusters frequently exhibit negligible interaction effects with top-tier classifiers, suggesting that performance gains are often dominated by the individual efficacy of the components rather than by synergistic pairings. This should reassure data analysts that prioritizing the individual quality of adjusters and classifiers is a robust heuristic, as the inherent strengths of top-tier algorithms often mitigate the necessity for case-specific pairings.

### Warnings

Imbalanced data can cause heavy confounding between the batch and target variables. Class differences are expected to induce differential expression—class imbalance often manifests as a mean shift that cannot be easily isolated from technical batch effects. Adjusters that are unaware of the class imbalance can inappropriately merge disparate groups. Using class labels or known groups for batch adjustment—supervised adjustment—addresses this problem, but can lead to poor generalization through model misspecification or overfit. Figure 4 demonstrates this effect. Supervised ComBat correction generally performs worse than the unsupervised version due to improper class effect assumptions. However, when training datasets are highly imbalanced, the assumptions of unsupervised ComBat (equal populations, a common generating distribution) are miscalibrated. All unsupervised adjustments tested here exhibit poor performance when training sets are highly imbalanced. In these situations, supervised adjustments can control for the confounding effects. However, this benefit may only be seen in the most pronounced imbalance scenarios.

**Figure 4:**
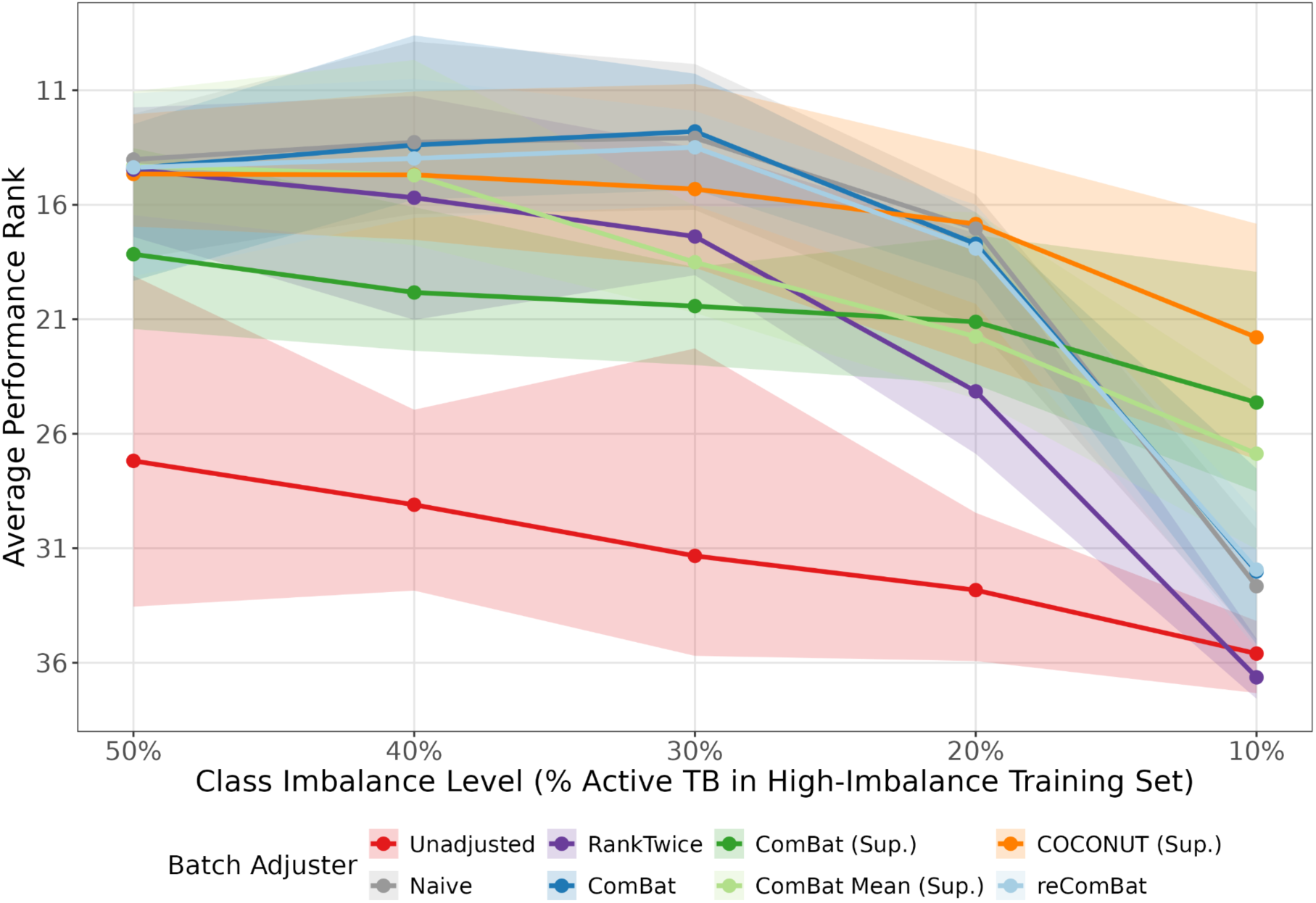
Batch Correction Performance using Training Sets with Opposite Label Imbalances. Colored lines compare various supervised and unsupervised adjustments. The y axis shows mean rank comparing the strategies for each test set and iteration, and imbalance level. Class imbalance level increases from left to right. A class imbalance of 20% indicates that one training dataset was filtered to have 20% active samples, while the second was filtered to have 80% active samples. Sample sizes were held constant over imbalance levels. Ranks were averaged over the two top performing classifiers (ElasticNet and XGBoost). Shaded region shows the minimum and maximum performance over test sets and training pairs.

### The Limitations of Internal Cross-Validation

Internal validation should not be substituted for cross-study prediction, as internal metrics often fail to capture real-world data variability. In k-fold cross-validation—a method frequently preferred over standard train-test splitting—the merged data is partitioned into k distinct validation groups [74]. The classifier is iteratively trained using data from k-1 groups and evaluated on the remaining held-out group until every sample has been tested. While this approach is rigorous for internal assessment, it often produces optimistic performance estimates because the model can learn batch-specific patterns that do not generalize [75]. This limitation is evidenced in Figure 5, while independent test studies show significant performance deltas. Because internal validation groups represent unseen samples but fail to simulate the unique variability of unseen batches, cross-batch testing on independent validation cohorts is essential to account for batch effects and obtain realistic performance benchmarks.

**Figure 5:**
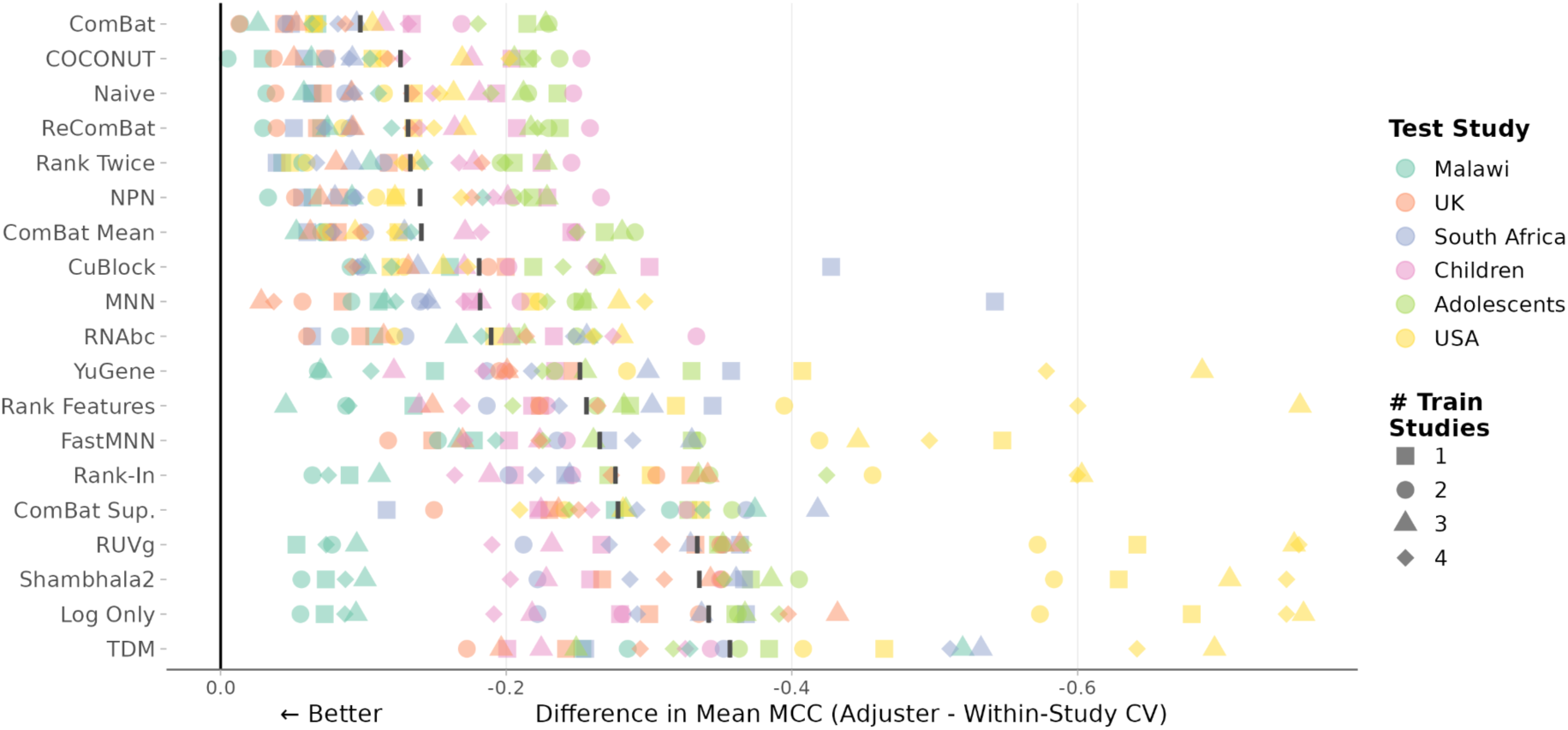
Adjuster Performance vs. Within-Study CV. This figure compares the distribution of cross-study MCC to within-study 3-fold cross-validation MCC across the adjusters. Points represent a particular test set (color), trained with a number of training datasets (shape), averaged over classifiers. Datasets were adjusted using the adjusters on the y axis. Cross-validation results are represented by the 0 baseline. All adjusted cross-dataset results show a noticeable gap from internal cross-validation performance. Internal cross-validation methods tend to provide optimistic performance ceilings for generalization to new batches.

### Batch Adjustment, Ensembling, and Meta-Analysis

Researchers seeking patterns across studies can choose between batch adjustment (data aggregation), ensembling (model and prediction aggregation), or meta-analysis (statistical aggregation) [76]. When batch adjustment is hindered by heavy confounding or severe heterogeneity, ensembling and meta-analysis provide viable alternatives that avoid direct data merging. Ensembling occupies a middle ground by training separate models on individual datasets and aggregating their parameters or outputs. This approach rewards features with stable predictive power across batches, providing a robust alternative to merging when datasets are highly heterogeneous [51]. Meta-analysis bypasses any knowledge of the raw data by combining statistical results. This approach is recommended when individual-level data used for the analysis are unavailable (potentially due to privacy constraints) or if the goal is to identify only the most robust, consistently replicated findings across studies [77]. For classifier development specifically, merging or ensembling are preferred because classifiers require access to sample-level data to learn decision boundaries. Meta-analysis of classifier performance across studies can complement this by assessing consistency but cannot replace the need for integrated training data.

### Generalizability to Other Omics

The integration of multi-study transcriptomic data holds immense potential for precision medicine, offering the statistical power and diversity necessary to train robust machine learning classifiers. The core principles discussed in this chapter extend across omics modalities. Systematic technical variation exists in all omics data, and often masquerades as biological signal. When features are jointly continuous, such as for bulk transcriptomics, proteomics, and metabolomics, location and scale adjustments are useful. Each modality may require specific adaptations, such as M-value transformations for DNA methylation or addressing extreme sparsity in metagenomics. When samples belong to clear groups, like cells sampled using scRNA, or imaged using Cell Painting, nearest neighbor approaches are applicable [13]. Even more complex data, such as spatial measurements, require graph-based or spatially aware correction methods that preserve local correlations [78]. The sensitivity of classifiers to batch effects is universal and necessitates the removal of technical noise while preserving biological signal. Care must always be taken to avoid overfitting when using target labels for correction. To ensure that molecular signatures represent genuine disease biology rather than dataset-specific noise, pipeline development must prioritize external, cross-study validation.

### Code and Data Availability

Source code is available at https://github.com/prestonraab/mitigation_impacts_on_classifiers. The pixi package manager [79] and Snakemake workflow management system [80] were used to ensure reproducibility.

### Use of Artificial Intelligence

During the preparation of this work, the author used Google Gemini to assist in drafting and refining the manuscript’s prose. After using this tool, the author thoroughly reviewed and edited the content and takes full responsibility for the accuracy and integrity of the final publication.

